# StrainPro – a highly accurate Metagenomic strain-level profiling tool

**DOI:** 10.1101/807149

**Authors:** Hsin-Nan Lin, Yaw-Ling Lin, Wen-Lian Hsu

## Abstract

Characterizing the taxonomic diversity of a microbial community is very important to understand the roles of microorganisms. Next generation sequencing (NGS) provides great potential for investigation of a microbial community and leads to Metagenomic studies. NGS generates DNA fragment sequences directly from microorganism samples, and it requires analysis tools to identify microbial species (or taxonomic composition) and estimate their relative abundance in the studied community. However, only a few tools could achieve strain-level identification and most tools estimate the microbial abundances simply according to the read counts. An evaluation study on metagenomic analysis tools concludes that the predicted abundance differed significantly from the true abundance. In this study, we present StrainPro, a novel metagenomic analysis tool which is highly accurate both at characterizing microorganisms at strain-level and estimating their relative abundances. A unique feature of StrainPro is it identifies representative sequence segments from reference genomes. We generate three simulated datasets using known strain sequences and another three simulated datasets using unknown strain sequences. We compare the performance of StrainPro with seven existing tools. The results show that StrainPro not only identifies metagenomes with high precision and recall, but it is also highly robust even when the metagenomes are not included in the reference database. Moreover, StrainPro estimates the relative abundance with high accuracy. We demonstrate that there is a strong positive linear relationship between observed and predicted abundances.

## Background

Microorganisms are extremely diverse and they play crucial roles in the different environments and human health. Some are a vital component of fertile soils; some have been used to convert carbohydrates to alcohol in the fermentation process; some colonize in human bodies and perform specific tasks useful to the human host; while some are pathogenic or harmful to human health. Characterizing the taxonomic diversity of a microbial community is very important to understand the roles of beneficial and harmful microorganisms. However, a majority of microorganisms can only survive in specific habitats and cannot be cultured in the laboratory[1]. Next generation sequencing (NGS) provides great potential for investigation of a microbial community and leads to Metagenomic studies. NGS generates DNA fragment sequences directly from microorganism samples, and it requires analysis tools to identify microbial species and estimate their abundance within the sampling community. A number of analysis tools have been developed, which can be categorized into two groups: those targeting the entire genomic content of a sample (i.e., metagenomics) and those focusing on marker genes (i.e., metataxonomics) [2]. The former group includes Centrifuge[3], CLARK[4], Genometa[5], GOTTCHA[6], Kraken[7], KrakenUniq[8], MEGAN[9], Sigma[10], etc. The latter group includes MetaPhlAn[1, 11], MetaPhyler[12], mOTU[13], QIIME[14], etc. Though metataxonomics provides information of species composition of a microbiome, the marker gene-based (e.g. 16S rRNA gene for bacteria) methodologies can only capture organisms that have the conserved genes, and it is estimated >50% of the organisms evaded detection with classical 16S amplicon sequencing [15]. More importantly, the marker gene-based methodologies are often insufficient for discrimination at the species and strain levels of classification [16, 17]. Shotgun based methodologies can overcome the limitations by targeting the entire genomic content of a sample. Moreover, the data obtained by shotgun approach are getting more complete and the sequencing cost is getting less expensive, the use of shotgun metagenomics is becoming more popular [18]. In this study, we focus on developing new approaches to analyze shotgun metagenomic data. Metagenomic analysis tools are developed to characterize the taxonomic composition of the studied community. They can be further classified into three classes [19]: 1) kmer-based; 2) alignment-based; and 3) Bayesian or EM-based classifiers. All these methods compare the sequencing data with reference databases. Among these methods, only a few methods could achieve strain-level identification. Strain or sub-species is a low-level taxonomic rank to differentiate microorganisms of the same species. They are more important to clinical diagnostics or pathogen identification. For example, most Escherichia coli strains in the human intestine are non-pathogenic, while only a certain strains of E.coli can cause disease [20]. O157:H7 group is one example of E.coli strains that produces potentially lethal toxins. Therefore, strain-level taxonomic assignment of a metagenomics-based study can provide more useful information in clinical diagnostics or epidemiological tracking.

However, to achieve strain-level identification, an analysis tool requires a comprehensive database of reference genomic sequences, which can result in a huge amount of sequence comparison and therefore become a bottleneck in the metagenomic data analysis. For example, the RefSeq database contains around 16,500 complete bacterial genomes in 2019, which consists of 55 billion nucleobases. Particularly, metagenomic data often involves hundreds of millions of read sequences. It requires an efficient algorithm to overcome all these challenges.

Another important issue is the estimation of relative microbial abundance, which is critical for understanding the microbial ecology of the environment and human health [21]. Most metagenomic analysis tools estimate the microbial abundances simply according to read counts. However, the estimation could be biased dramatically due to the genome size and sequence uniqueness. An evaluation study [2] on metagenomic analysis tools concluded that the predicted abundance differed significantly from the true abundance, and there was a large variation in the predicted abundance estimations of phyla between tools.

Here we briefly describe a few representative algorithms of metagenomic data analysis tools. CLARK collects all unique k-mers in the reference genomes and it classifies a read to a reference genome if they share the highest number of k-mers. Centrifuge reduces the size of reference genomes by compressing genomic sequences and builds a FM-index for those compressed sequences. Firstly, it compares two highly similar genomes, say G_1_ and G_2_, using a genome comparison tool and discards sequences of G_2_ that are ≥99% identical to G_1_. The unique sequences of G_2_ are then added to G_1_, forming a merged genome, G_1+2_. This process repeats for the entire database until no merged genomes have sequences ≥99% identical to any other genome. Centrifuge then maps reads against the FM-index and assigns each read a single or multiple taxonomic labels. Kraken builds a database of k-mers from reference genomes and assigns a taxonomic identifier (taxid) to each k-mer according to the LCA (lowest common ancestor) of all organisms whose genomes contain that k-mer. It classifies reads by querying the database using the k-mers in read sequences. The taxa associated with a read’s k-mers form a taxonomy tree and the classification is made by finding the leaf node with the highest score in the tree. MetaPhlAn2 uses a set of around one million markers to detect the taxonomic clades present in a microbiome sample. The markers are substrings of reference genomes that represent clade-specific sequences. It then uses Bowtie2 to map all reads against the marker set and assign taxid accordingly.

In this study, we present StrainPro, a novel metagenomic analysis method, which is highly accurate both at characterizing microorganisms at strain-level and estimating their relative abundance. A unique feature of StrainPro is it identifies representative sequence segments from reference genomes. A representative sequence segment is a DNA substring that can be only found in genomes of a specific taxon and can be considered a signature for that taxon. Thus identical DNA fragments can be collapsed into a single copy. StrainPro searches NGS short reads against the representative sequence database and characterizes the underlying taxonomic composition of the metagenomic sample. The experiment result shows that StrainPro can identify strains and estimate their relative abundances more accurately than existing tools. We implemented StrainPro in C/C++, and the source codes and benchmark datasets are available at https://github.com/hsinnan75/StrainPro.

## Methods

StrainPro is an alignment-based method. It contains three major components: a) representative sequence identification; b) read mapping; and c) strain composition identification. We describe the detailed algorithm of each major component below.

### Finding representative sequence segments

To compile a comprehensive database of reference genomic sequences, we download all completed microbial genomes in the NCBI RefSeq database. We then cluster those genomes according to taxonomic information. At first, genomes of the same species are clustered together. We define the cluster size as the total genome sizes in that cluster. If a cluster size is less than one billion nucleobases, we then merge those clusters with the same genus (or family, order, and so on) until the merged cluster meets the threshold or there are no more such clusters. Since each taxon of a particular ranking has different number of genomes, not every cluster is formed by gathering genomes with the same taxonomic rank. For example, a cluster may only contain genomes of Escherichia coli (rank: species) since E.coli is a large and diverse group, while another cluster contains genomes of Proteobacteria (rank: phylum) since the underlying sub-groups are not large enough to form an independent cluster. After clustering, we build a BWT index for each genome cluster. Thus, we will build *n* BWT indexes if there are *n* genome clusters.

Since genomes are clustered based on taxonomic information, those which belong to the same taxid share a certain degree of sequence similarity and each of the genomes may also possess a certain degree of sequence uniqueness. Thus we can divide genome sequences into different segments with variable length and assign each to the LCA according to the taxids of involved genomes. For example, a sequence segment is assigned to 83334 (taxid of E.coli O157:H7 strains) if it can only be found in those strains; and another segment is assigned to 1224 (taxid of Proteobacteria) if it can be found in diverse genomes whose LCA is 1224. Once we collect all representative sequence segments from the original genome database, we can build a more compact database for read classification.

To identify the representative sequence segments, we introduce the idea of pseudo-reads. Given a reference genome *G* of size *N*, let *G*_*i*_ be the *i*-th nucleobase of *G* and *G*[*i, j*] be the substring between *G*_*i*_ and *G*_*j*_. We define ps_read_*i*_ is a pseudo-read of length *l* whose sequence is *G*[*i, i+l*-1]. Thus we have *N*-*l*+1 pseudo-reads in *G* and each read is used to search against the cluster that contains *G*. Suppose ps_read_*i*_ can only be found in *G*, we assign *G*_*i*_ to the taxid of *G*; otherwise we assign *G*_*i*_ to the LCA taxid of the genomes containing that pseudo-read. Notably, if ps_read_*i*_ has multiple occurrences in *G* or in other genomes, we only assign the first occurrence to the LCA taxid, and the others are assigned to 0 meaning they are redundant. After all pseudo-reads have been checked for every genomes in that cluster, we convert each genome sequence into a long list of taxids and 0s. We perform a linear scan through each long list and divide it into sub-lists if any two adjacent taxids on the list are different. Thus each sub-list indicates a representative sequence segment associated with the same taxid. We ignore all sub-lists which are associated with 0. Notably, the pseudo-read length *l* is a trade-off between specificity and sensitivity. A small *l* will lose the specificity of a pseudo-read though it is more likely to have more occurrences among reference genomes; A large *l* will lose the sensitivity of a pseudo-read though it is more likely to capture the uniqueness of a strain sequence. By default StrainPro uses 101-bp since it is the most common length of NGS short reads. The user can modify the value based on their NGS data.

Figure 1 shows an example illustrating our idea of representative sequence segments. In this example, there are two genomes in the cluster, G^1^(taxid:A) and G^2^(taxid:B), and we assume LCA(A,B) = C. Each pseudo-read is used to search against the two genomes and is assigned to taxid of A, B, C, or 0 according to its occurrences. The search result shows that G^1^ can be divided into five segments which are associated with taxid of A, C, A, C, A respectively, and G^2^ can be divided into two segments which are both associated with taxid of B. Those segments which are associated with 0 are redundant sequences and they are discarded in the compact database construction.

**Figure 1.**
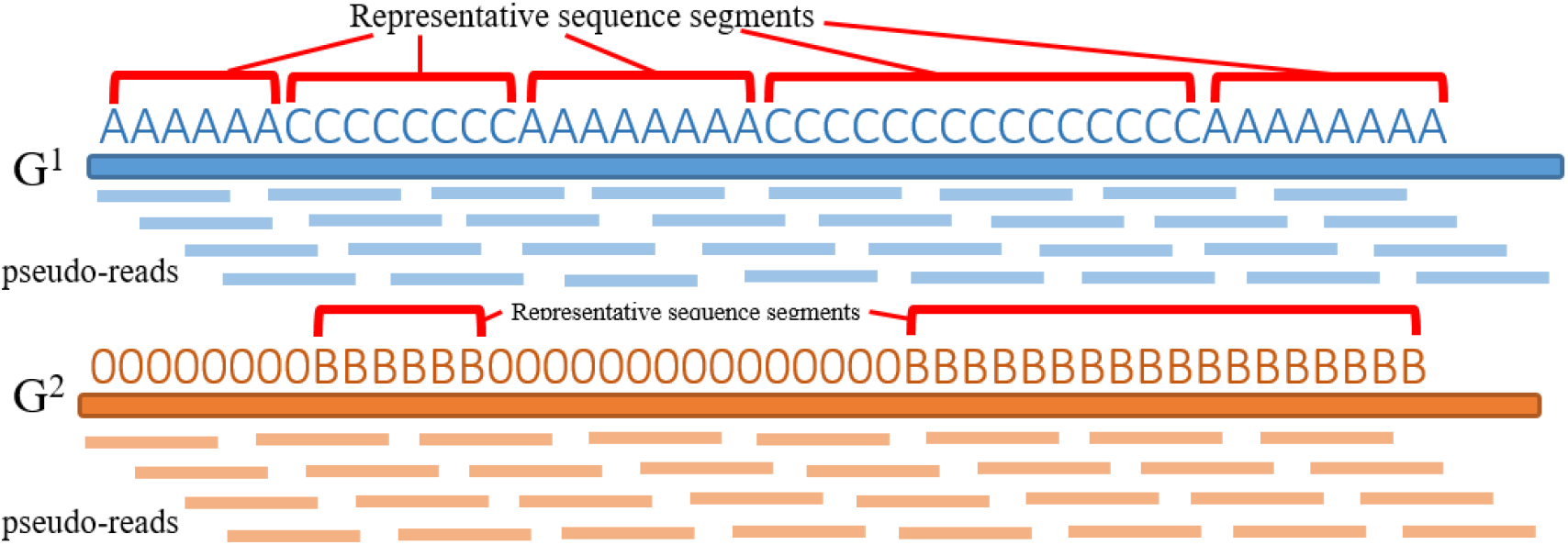
The identification of representative sequence segments with two genomes. G^1^(taxid:A) and G^2^(taxid:B), and LCA(A,B) = C.

We replace the original reference genomes with those representative sequence segments and cluster those segments the same way we do for the original reference genomes. Notably, each representative sequence segment is considered as an independent sequence entry. The resulting clusters are used to build BWT indexes as the compact database for read mapping.

### Read mapping

Since the representative sequences are clustered into multiple groups according to the taxonomic information, we build a BWT index for each representative sequence cluster. StrainPro uses a modified algorithm of KART to perform read mapping. The detailed read mapping algorithm of KART can be found in our previous study [22]. Here we focus on the high-level methodology description. StrainPro adopts a divide-and-conquer strategy to handle matches and mismatches separately between read sequence and reference genome. StrainPro identifies all locally maximal exact matches (LMEMs). We then cluster those LMEMs according to their coordinates and fill gaps between LMEMs to create candidate alignments.

StrainPro aligns every individual read sequence onto each representative sequence cluster and assigns it to the taxid of the mapped representative sequence segment with the highest alignment score. If a read has multiple best alignments, it is assigned to the LCA of those hits. Although we have removed redundant sequences by using pseudo-reads, a read could still have multiple hits due to sequence variations or the duplications locating in different clusters. The read mapping is performed iteratively until all the representative sequence clusters have been searched.

### Identifying taxonomic composition and estimating relative abundance

Each representative sequence segment associated with a taxid is a basic unit for identifying the taxonomic composition and estimating relative abundances. We infer the presence and absence of a taxid by quantifying the reads that are mapped onto a representative sequence segment. Thus we can differentiate between true alignments and false alignments by checking the read density in the corresponding representative sequence segment.

To explain the detailed implementation of this concept, we here use a 3-tuple, (*Taxid*_*s*_, *Freq*_*s*_, *Size*_*s*_) to denote the mapping profile of a representative sequence segment *S*, where *Taxid*_*s*_ is the taxid of *S, Freq*_*s*_ is the number of reads that are mapped onto *S*, and *Size*_*s*_ is number of all pseudo-reads of *S*. It can be expected that if a genome containing *S* is present in the metagenomic sample, *Freq*_*s*_ should be in direct proportion to the abundance of *S* theoretically and empirically. Furthermore, a sequencing depth of *d* implies that the probability of a short read of length *l* starting at a specific position is approximately equal to *d / l*. Therefore, the ratio *R*_*s*_ of *Freq*_*s*_ to *Size*_*s*_ is positively correlated with *d / l*. Based on this theoretical model, we can calculate the sequencing depth of each strain and estimate relative abundance based on their sequencing depths. We here use an example to demonstrate how we infer taxonomic composition with representative sequence segments. Suppose we are given two mapping profiles of representative sequence segments *A* and *B*, which are (*Taxid*_*A*_, 100, 100) and (*Taxid*_*B*_, 100, 1000). Although both the segments contain 100 of read count, the mapping density of segments *A* is much larger than that of segment *B*. It implies that reads that are mapped onto *B* are more like random mappings. In such cases, segment *B* is less likely present in the metagenomic sample.

To prevent false identification of taxonomic composition, we set a threshold *t* on the ratio *R*_*s*_. We only consider *S* to be present in the metagenomic sample if *R*_*s*_ > *t*. Thus given a *taxid*_*s*_ at strain-level, the sequencing depth is estimated as follows:

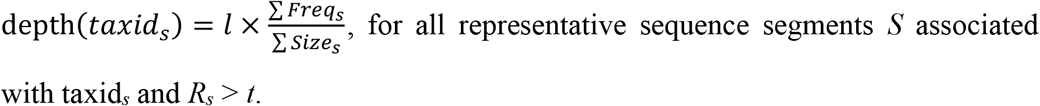

The relative abundance of taxid_*s*_ is then estimated as follows:

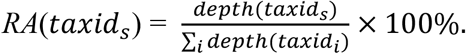

To estimate the relative abundance of a taxid at species-level and above, we adopt an alternative metric since genomes between different species are more distinguishable and not every microorganism is classified into strain-level. If we only use the abundance of taxids at strain-level to infer those at species-level, the estimation may not cover the whole present microorganisms in the metagenomic sample. Thus for those taxids at species-level and above (denoted as *taxid*_*p*_), we estimate their abundance using the number of reads that are mapped onto a representative sequence segment *P*. Notably, *P* here represents all representative sequence segments that are associated with *taxid*_*p*_ or its sub-types and *R*_*p*_ > *t*. The relative abundance of *taxid*_*p*_ is estimated as follows:

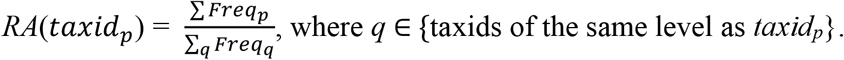

## Results

### Simulated metagenomic datasets and performance metrics

The genuine metagenomic samples lack ground-truth taxonomic compositions. The true strains and species in those samples are mostly unknown. It is difficult to use genuine metagenomic datasets to assess performance on the taxonomic predictions in this study. Instead, we generate simulated taxonomic compositions using real strains in NCBI RefSeq database. The NCBI RefSeq database contains around 16,500 complete bacterial genomes, where 6,200 genomes are associated with taxids of strain-level and the other 10,300 genomes are with species-level. In this study, we generate three simulated taxonomic compositions which each contains around 50 random strain-level genomes. Since strains are randomly chosen from the 6,200 genomes, some of them may belong to the same sub-species. We use the whole database to generate the representative sequence segments. For each simulated taxonomic composition, we use the WGSIM program (https://github.com/lh3/wgsim) to generate metagenomic reads with read length 101bp, 0.2% sequencing error rate and 50X sequencing depth. The three simulated metagenomic datasets are referred to as SimData_1, SimData_2, and SimData_3 respectively. To assess the taxonomic prediction accuracy on unknown strains, we also simulate novel strains by modifying genome sequences in each simulated taxonomic composition. We use WGSIM to alter genome sequence with 1% of mutation rate (15% of which are INDELs and 85% are SNPs) to simulate polymorphism for novel strains and generate their metagenomic reads with the same arguments. The three simulated metagenomes of novel strains are referred to as SimNovel_1, SimNovel_2, and SimNovel_3 respectively. Table 1 shows the strains in the three simulated taxonomic compositions.

**Table 1.**
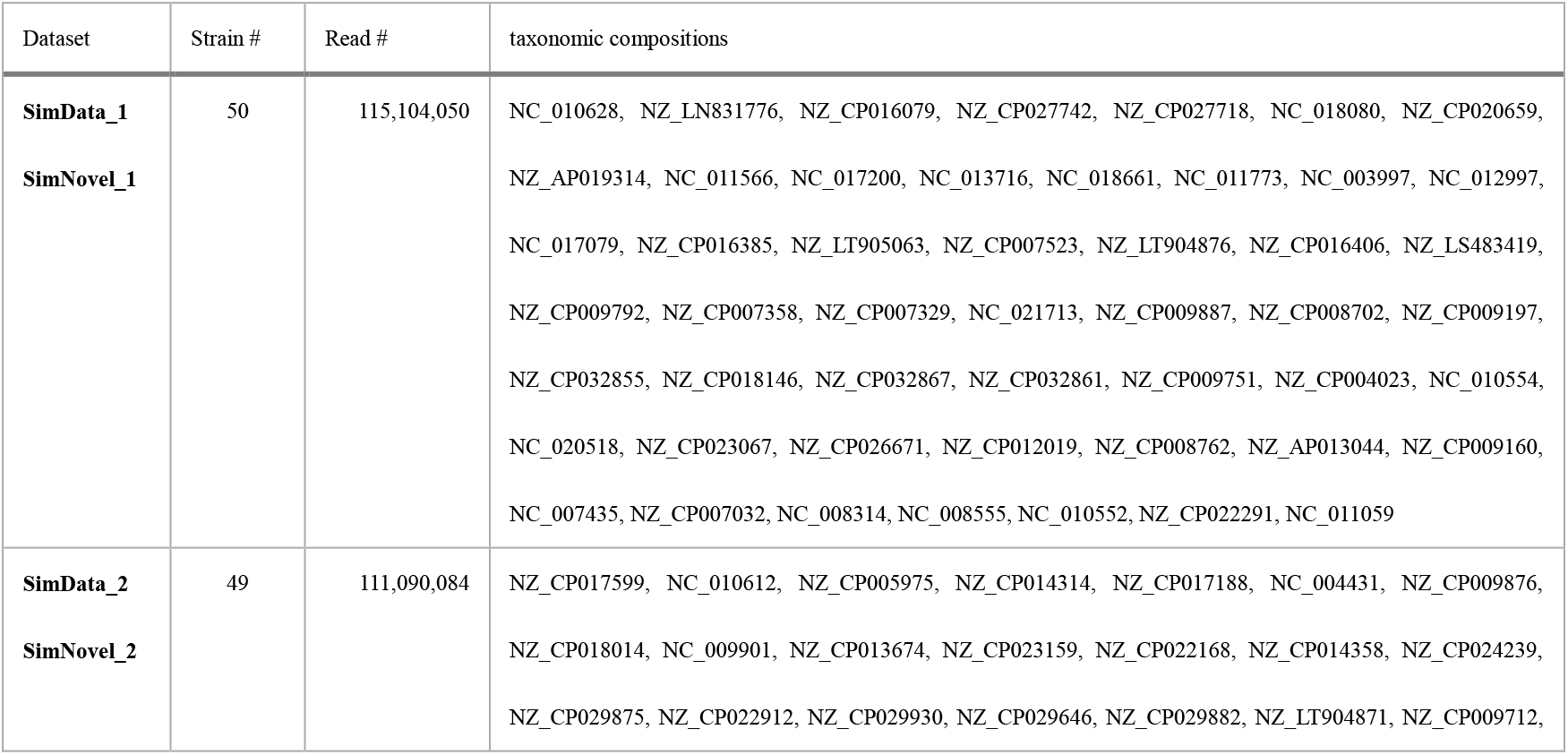
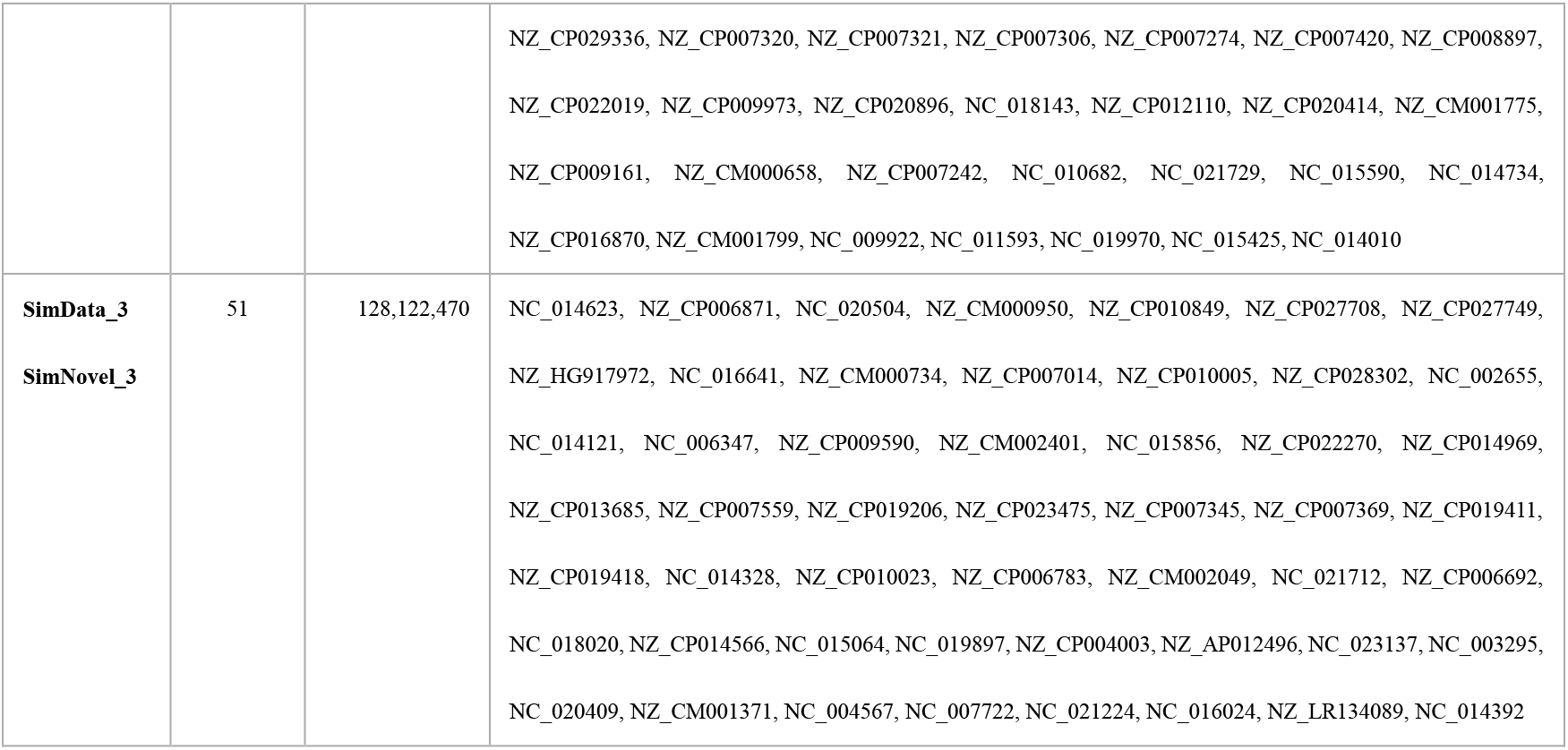
The taxonomic compositions and number of reads in the simulated datasets. The genome sequences in SimNovel_1/2/3 are modified with 1% of mutation rate to simulate polymorphism for novel strains.

We use precision, recall, and run-time to evaluate the performance of each tool. A predicted taxid is considered a true positive (TP) if it is present in the simulated metagenome and its read count above a certain threshold; a predicted taxid is considered a false positive (FP) if it is absent in the simulated metagenome and its read count above the threshold. Given a simulated metagenome of *N* strains, precision is defined as TPs / (TPs+FPs) and recall is defined as TPs / N.

In this study, we compare the performance of StrainPro with seven existing tools, including Centrifuge, CLARK, GOTTCHA, Kraken2, KrakenUniq, MetaPhlAn2, and Sigma. All these tools are able to achieve strain-level classification except CLARK and Sigma. We tried to include more other metagenomic analysis tools; however, some tools failed in the middle of execution, some lacked reference database for all microorganisms, and some were designed for marker genes only.

### Performance comparison on simulated metagenomes (original strains)

In this comparison, we assess the precisions and recalls of each method at strain-, species-, and genus-levels. Figure 2 illustrates the comparison result on the three simulated metagenomes. Notably, some tools only perform taxonomic classification for every read without generating the taxonomic composition. In such cases, we write a simple script to infer the taxonomic composition based on read count of each associated taxid. We also find a proper threshold for each tool to filter out some taxids according to their performance. The commands as well as the reference database and argument setting used for each tool are shown in Table S1 in the supplementary materials.

**Figure 2.**
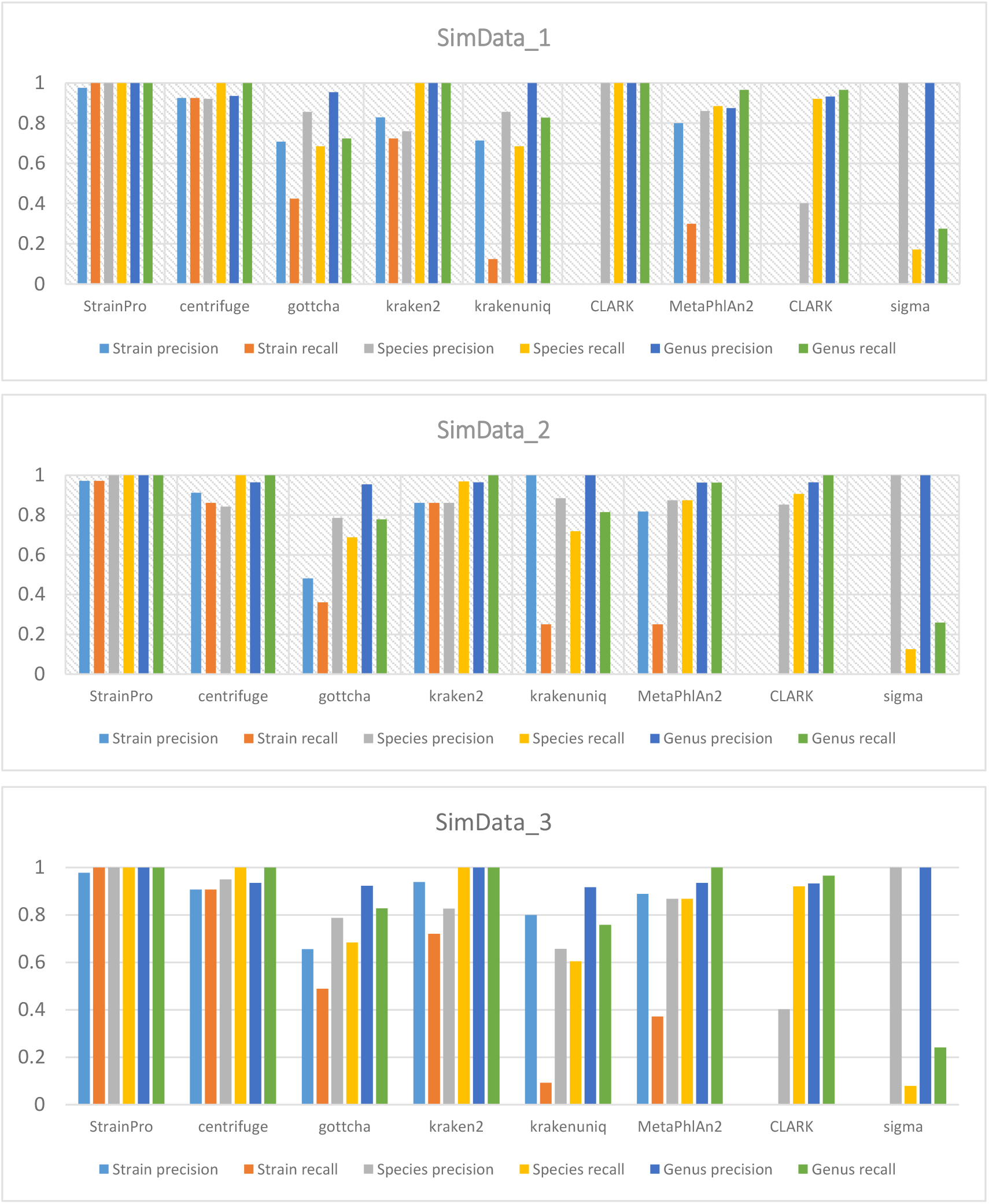
Performance comparison on the three simulated metagenomes (present strains are included in the reference database).

It can be observed that StrainPro achieves the best precisions and recalls at the three taxonomic levels among the selected tools. In particular, StrainPro consistently yields higher precision and recall at the strain-level than the other tools. For example, the F1-scores for StrainPro on the three datasets are 0.988, 0.972, and 0.989 respectively. Though CLARK is not able to achieve strain-level resolution, its precisions and recalls at the species- and genus-levels are very close to those of StrainPro. Centrifuge yields the second best performance in general, while Kraken2 performs slightly worse than Centrifuge. Centrifuge’s F1-scores are 0.925, 0.886, and 0.907 respectively. The F1-scores at strain-level for Kraken2 are 0.773, 0.861, and 0.816 respectively. KrakenUniq is derived from Kraken, however, its performance is surprisingly not as good as Kraken2. It yields much worse recalls than Kraken2. Its F1-scores at strain-level are 0.213, 0.400, and 0.167 respectively. MetaPhlAn2 produces good performance both at species- and genus-levels, but its recall at strain-level is not satisfactory. Likewise, GOTTCHA’s precisions are higher than its recalls. We assume their pre-built databases are not comprehensive enough to cover all kinds of strains and customized databases are not supported yet for both tools. Though Sigma uses the whole NCBI RefSeq as the reference genomes, there is a huge gap between its precision and recall at species- and genus-levels. It achieves perfect precisions, while all its recalls are around 25% for the three metagenomes.

### Performance comparison on simulated metagenomes (novel strains)

In this experiment, we simulate novel strains by altering the original strains with 1% of mutation rate. The purpose is to assess the prediction accuracy of a metagenomic analysis tool when the present genomes are not included in the reference database. In this comparison, we only assess the prediction accuracy at species- and genus levels since sequence modification has changed the original strain sequences, the original strain-level classification is not applicable. Figure 3 shows the comparison result on the three simulated metagenomes. Here we use the same thresholds to filter out taxids according to their read counts.

**Figure 3.**
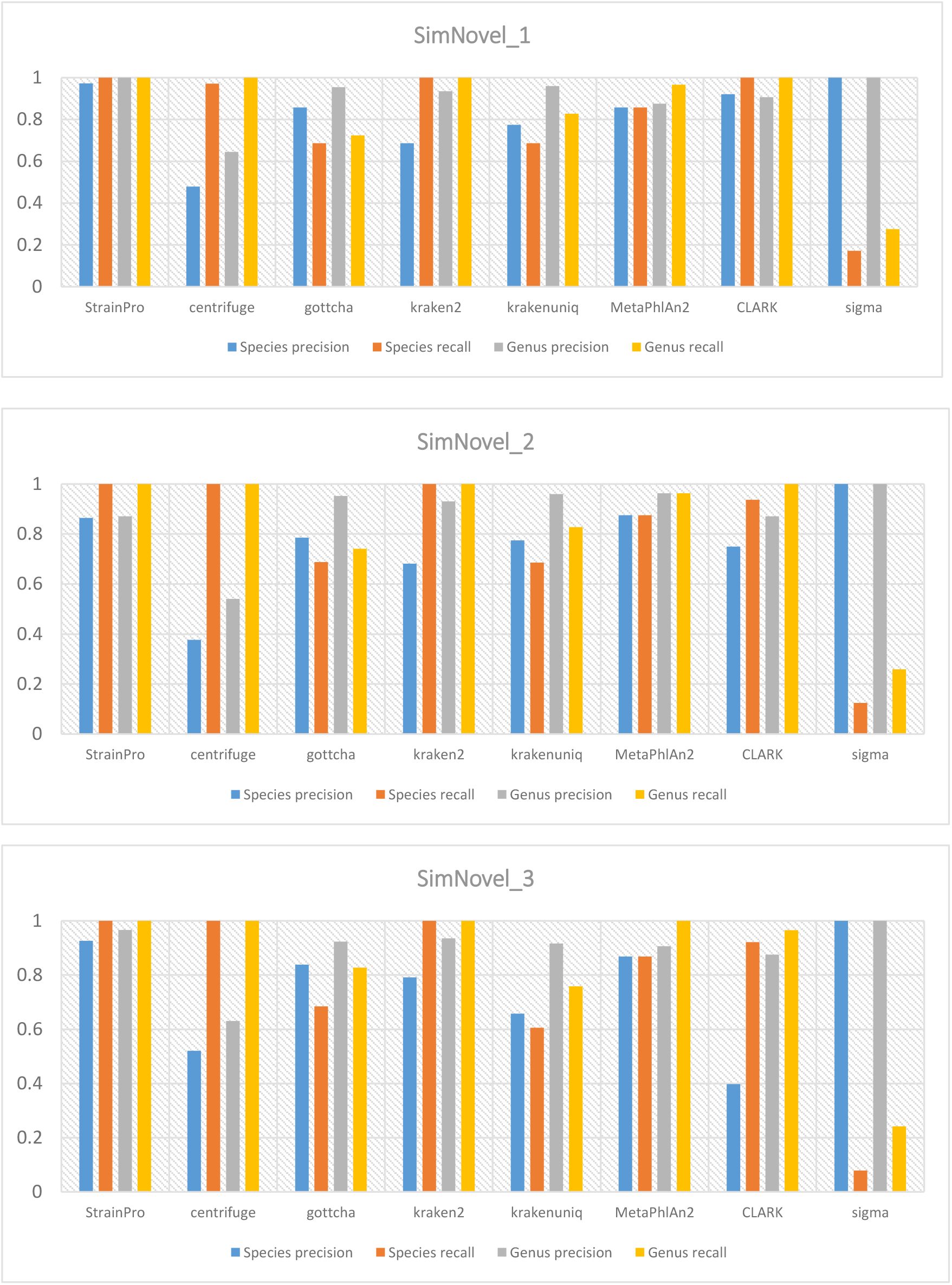
Performance comparison on the three simulated metagenomes (present strains are not included in the reference database).

It can be observed that StrainPro still produces the best performance on these metagenomes. We demonstrate that StrainPro is very robust even though the novel strains are not included in the reference database. StrainPro achieves the highest F1-scores at species- and genus-levels. Interestingly, Centrifuge is the second best tool in the previous experiment; however, its precision declines significantly with the same threshold of read count. We could have increased the threshold to improve its precisions, but the manipulation of threshold values is based on the knowledge of metagenomes. There are no clear rules on the threshold setting when it comes to real datasets. MetaPhlAn2 generally performs the second best on those metagenomes. It suggests MetaPhlAn2 is also robust to novel strains. Its precisions and recalls at species- and genus-levels are very similar to those on original metagenomes. CLARK and Kraken2 also produce comparable performance to MetaPhlAn2, though CLARK yields less satisfactory precision on SimNovel_3, and Kraken2’s precisions at species-level on mutant metagenomes are worse than those on original metagenomes. GOTTCHA performs comparably to KrakenUniq. Their performance is also similar to that on original metagenomes. Likewise, Sigma achieves perfect precisions, while its recalls are also around 25% for the three metagenomes.

### Performance comparison on classification speed

Due to the huge amount of short reads in metagenomic data and the size of reference databases, the classification speed is also an important issue. We assess the classification speed for each tool on the three original metagenomes. Each tool is evaluated its speed using 16 threads. Notably, the run-time for database creation is not considered here. Figure 4 illustrates the comparison result. It can be observed that Centrifuge, Kraken2, KrakenUniq, and CLARK perform similarly in term of classification speed. These tools are able to classify more than 100 million reads in a half hour or less. StrainPro, GOTTCHA, and MetaPhlAn2 are the second fastest groups. They spend around two or three hours on each metagenome. Sigma is the lowest tool in this comparison. It spends more than 10 hours on each metagenome.

**Figure 4.**
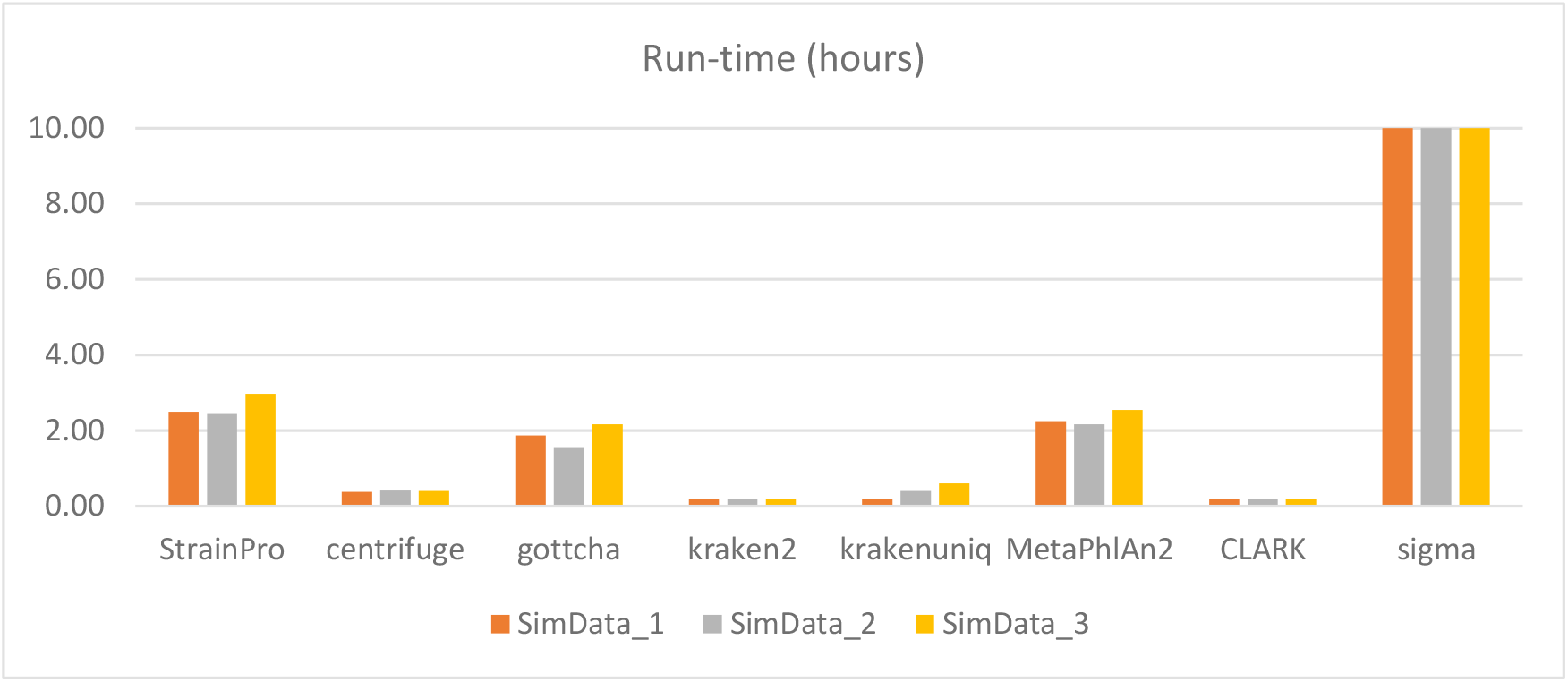
Run-time comparison on the three simulated metagenomes.

### The relative abundance estimation at strain-level

In this study, we estimate the relative abundance of each taxid based on the mapping density at representative sequence segments. It is very different from that simply based on read counts. We here analyze the average difference between the true and predicted abundance of each strain-level taxid. Given a strain-level taxid, if the true relative abundance is X%, and the predicted relative abundance is Y%, then the difference is defined as |X-Y|. We also measure the linear correlation between the two abundances using the Pearson correlation coefficient (PCC). Table 2 summaries the analysis result. In this analysis, we compare StrainPro to Centrifuge and Kraken2 since the three tools produce at least 0.8 of recall at strain-level prediction. It can be observed that StrainPro yields the smallest average difference between true and predicted abundances. The PCC scores of StrainPro are around 0.95. In contrast, Centrifuge and Kraken2 yield much larger average difference and their PCC scores suggest that there is no linear correlation between true and predicted abundances.

**Table 2.** The evaluation of relative abundance prediction in StrainPro, Centrifuge, and Kraken2.

| Dataset | Tools | Average difference | Pearson correlation coefficient |
| --- | --- | --- | --- |
| SimData_2 | StrainPro | **0.29** | **0.943** |
|  | Centrifuge | 2.89 | -0.083 |
|  | Kraken2 | 2.48 | 0.176 |
| SimData_3 | StrainPro | **0.68** | **0.967** |
|  | Centrifuge | 3.79 | -0.069 |
|  | Kraken2 | 2.51 | 0.444 |
| SimData_4 | StrainPro | **0.20** | **0.986** |
|  | Centrifuge | 2.56 | -0.046 |
|  | Kraken2 | 2.76 | -0.013 |

### Representative sequence segment analysis

The identification of representative sequence segments is a unique design in this study. We demonstrate the representative sequence segments are more suitable to classify short reads and identify the metagenomic composition. We here analyze the efficiency of representative sequence segments.

The RefSeq database contains 16,500 complete bacterial which are separated into 28 clusters according to our algorithm. We then identify representative sequence segments in each cluster. The original cluster size and the resulting representative sequence segment size are shown in Figure S1 in the supplementary materials. The original reference genomes contain 54.94 billion nucleobases. After identifying the representative sequence segments and removing redundant sequence segments, the remaining sequences only contain 26.96 billion nucleobases. We reduce the database size by around 50%. The more strains of the same species in a cluster, the more redundant sequences we can remove. For example, the original cluster size of E.coli strains is 3.6 GB, and the resulting representative sequence’s size is only 0.29 GB. We reduce the reference genome size of E.coli strains by more than 90%. However, we still keep all the unique sequence fragments of each E.coli strain.

## Conclusions

In this study, we describe a novel metagenomic analysis tool, called StrainPro to classify NGS short reads and identify the underlying metagenomic composition at strain-level and above. We define representative sequence segments as the sequence features for each taxonomic identifier. The relative abundance estimation is based on the mapping density at the representative sequence segments. Thus StrainPro is able to differentiate between true mapping and false mapping by measuring the mapping density in the representative sequence segments. We demonstrate the mapping density is useful to filter random alignments out. Moreover, identifying the representative sequence segments can also remove redundant sequences and reduce the database size significantly.

To assess the classification accuracy, we generate three simulated metagenomes using known strain sequences and another three simulated metagenomes using mutant strain sequences. We compare the performance of StrainPro to seven existing tools. The comparison results show that StrainPro not only identifies the metagenomic composition with high precision and recall, but it is also highly robust when the metagenomes are not included in the reference database. Furthermore, StrainPro estimates the relative abundance with high accuracy. The average difference between the true and predicted abundance is very small and the two abundances shows a highly positive linear relationship.

Though StrainPro is not as fast as some existing tools, it is able to identify strains and estimate their relative abundances more accurately than existing tools. More and more medical and biological researches adopt shotgun metagenomics to study taxonomic diversity of a microbial community nowadays, we are confident that StrainPro is able to provide reliable and accurate metagenomic data analysis.

