## Supplementary for "StrainPro – a highly accurate Metagenomic strain-level profiling tool"

Hsin-Nan Lin and Wen-Lian Hsu*
Institute of Information Science, Academia Sinica, Taipei, Taiwan

### Supplementary material

Table S1. The metagenomic data analysis tools and their commands and thresholds used for the benchmark datasets.

| Tool | Version | Arguments | Read count threshold |
|---|---|---|---|
| StrainPro | 0.9.0 | StrainPro-map -t 16 -i bacteria_idx -f read.fq -o res.txt | 50 |
| Centrifuge | 1.0.4 | centrifuge -p 16 -x bacteria_idx -q read.fq --report-file res.map > /dev/null | 1000 |
| GOTTCHA | 1.0c | gottcha.pl --threads 16 --outDir out --input read.fq --database database/GOTTCHA_BACTERIA_c4937_k24_u30.strain | 10000 |
| Kraken2 | 2.0.8-beta | kraken2 --threads 16 --db bacteria_db read.fq > read.map | 10000 |
| KrakenUniq | 0.5.8 | krakenuniq --threads 16 --db bacteria_db --report-file res.tsv --fastq-input read.fq > /dev/null | 10000 |
| MetaPhlAn2 | 2.7.7 | python metaphlan2.py read.fq --input_type fastq --nproc 16 > res.txt | N/A |
| CLARK | 1.2.6.1 | classify_metagenome.sh -n 16 -O read.fq -R res.txt | 10000 |
| Sigma | 1.0.1 (Beta) | 1. sigma-align-reads -p 1 -c dataset/config.cfg -w dataset (read mapping with 16 threads)<br>2. sigma -t 16 -c dataset/config.cfg -w dataset | N/A |

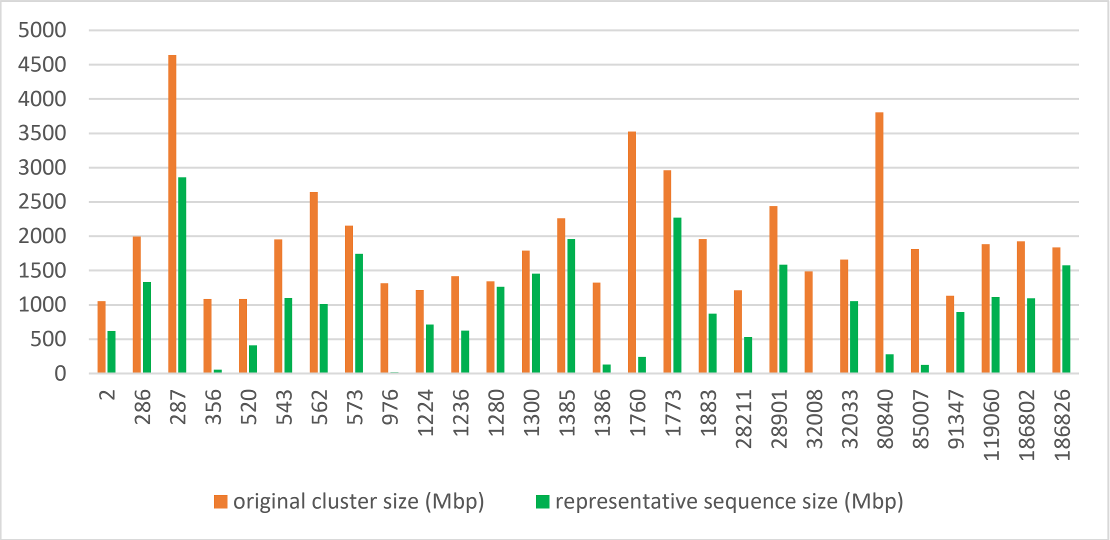


Figure S1. The original cluster size and the resulting representative sequence segment size. The x-axis indicates the taxid of each cluster. The y-axis indicates the cluster size in Mbp.
